# PHISDetector: a tool to detect diverse *in silico* phage-host interaction signals for virome studies

**DOI:** 10.1101/661074

**Authors:** Fan Zhang, Fengxia Zhou, Rui Gan, Chunyan Ren, Yuqiang Jia, Ling Yu, Zhiwei Huang

**Affiliations:** Harbin Institute of Technology; Harvard Medical School

## Abstract

Phage-microbe interactions not only are appealing systems to study coevolution but also have been increasingly emphasized due to their roles in human health, diseases, and novel therapeutic development. Meanwhile, their interactions leave diverse signals in bacterial and phage genomic sequences, defined as phage-host interaction signals (PHISs), such as sequence composition, CRISPR targeting, prophage, and protein-protein interaction signals. We infer that proper detection and integration of these diverse PHISs will allow us to predict phage-host interactions. Here, we developed PHISDetector, a novel tool to predict phage-host interactions by detecting and integrating diverse *in silico* PHISs and scoring the probability of phage-host interactions using machine-learning models based on PHIS features. PHISDetector is available as a one-stop web service version for general users to study individual inputs. A stand-alone software version is also provided to process massive phage contigs from virome studies. PHISDetector is freely available at http://www.microbiome-bigdata.com/PHISDetector/ and https://github.com/HIT-ImmunologyLab/PHISDector.

## Background

Phages play key roles in shaping the community structure of human and environmental microbiota and provide potential tools for precise manipulation of specific microbes. Recent studies have further revealed that phage-microbe interactions influence mammalian health and disease ^1^. Due to the great potential in developing novel therapeutics, such as phage therapy to combat multidrug-resistant infections, it is critical to identify and fully understand these interactions. Molecular and ecological coevolutionary processes of phages and microbes leave various signals in their genomic sequences to trace phage-host interactions ^2^. In addition to experimental methods, recent advances in large-scale genomic and metagenomic sequencing efforts and computational approaches have profoundly deepened our understanding of phage-microbe interactions and advanced new challenges in investigating such **<u>p</u>**hage-**<u>h</u>**ost **<u>i</u>**nteraction **<u>s</u>**ignals (PHISs).

PHISs can be grouped into mainly five categories and detected accordingly. First, PHISs can be detected by identifying putative prophage regions in bacterial genomes, defined as integrated phages that insert their genomes into their bacterial hosts. Several *in silico* tools for prophage detection in sequenced genomes have been developed, such as VirSorter ^3^, PHASTER ^4^, Prophinder ^5^, Phage_Finder ^6^, and PhageWeb ^7^. Recently, a microbe-phage interaction database (MVP) was developed mainly based on prophage inference ^8^. Second, based on the observation that phages share highly similar genomic signatures (such as *k*-mer or codon usage) with their hosts because phage replication is dependent on the translational machinery of its bacterial host ^9^, sequence composition analysis is a commonly used alignment-free method for PHIS detection. VirHostMatcher ^10^ and WIsH ^11^ are two tools developed to predict hosts for virus genomes or even short viral contigs based on *k*-mer signals. Third, CRISPR spacer sequences can also be used to infer phage-host interactions given that bacterial hosts incorporate spacer sequences from the phages that infect them ^12-14^. Fourth, genetic homology analysis based on homology between phage and bacterial genes is also used to identify phage-bacterial relationships ^15-17^. Fifth, protein-protein interactions (PPIs) have been applied to predict phage-host interactions because the interactions between a phage and a microbe are dependent mainly on the interactions between their encoded proteins ^18^.

Although various methods have been proposed to predict phage-host interactions using merely a single in silico signal, prediction accuracy and coverage is a limitation^2^. Meanwhile, with the exponentially increasing number of viruses uncovered in virome studies, there is a massive demand for a tool that is capable of incorporating all types of PHISs and conveniently predicting the microbial hosts of viruses, especially for virome studies. However, to the best of our knowledge, all currently available tools are limited to certain interacting features, and there is no published web server implementation or informed stand-alone software available to integrate all types of PHISs for comprehensive prediction of global phage-host interactions. Here, we developed PHISDetector, a novel integrative tool to predict phage-host interactions by detecting and integrating diverse in silico PHISs, including sequence composition, CRISPR targeting, prophage, and protein-protein interaction signals, and scoring the probability of phage-host interactions using machine-learning models based on PHIS features. PHISDetector captures phage-host associations in a data-driven manner, and is presented as a software pipeline for phage-host interaction identification, annotation and analysis in a comprehensive and user-friendly manner (Fig. 1). PHISDetector can be run either as a web server (http://www.microbiome-bigdata.com/PHISDetector/) or as a stand-alone version on a standard desktop computer.

**Fig. 1.**
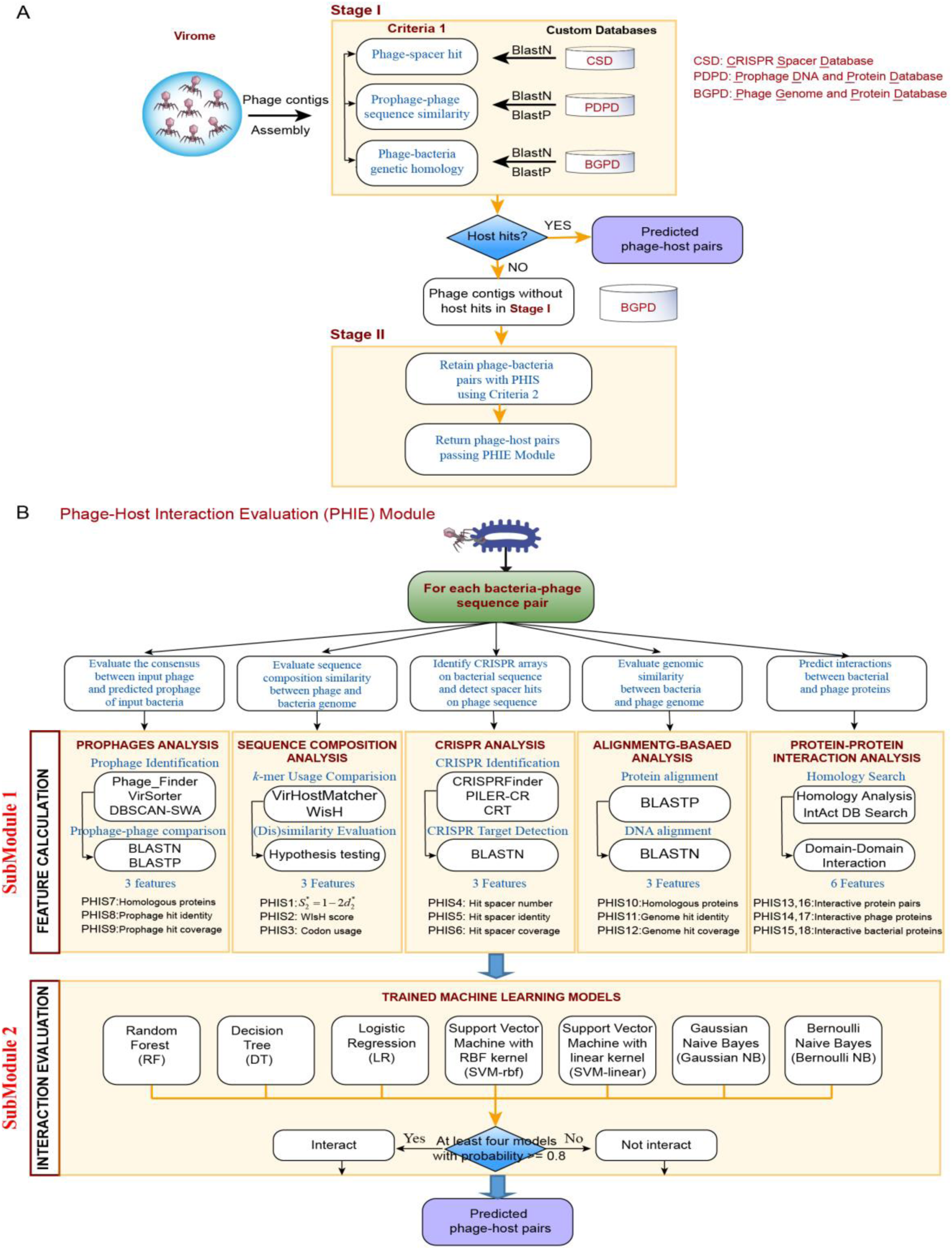
PHISDetector pipeline for prediction and evaluation of microbe-phage interactions. (a) PHISDetector combines two stages. In stage I, phage-host pairs with high reliability are detected using criterion 1 and returned directly as final predicted results. In stage II, phage-host pairs with potential PHISs based on criterion 2 are retained and further evaluated using trained machine-learning models implemented in the PHIE module. (b) Eighteen features belonging to five categories were calculated using sequence composition similarity, CRISPR targeting, prophage, genetic homology, and PPI or DDI analysis. Seven machine-learning models were then trained on the training dataset with these 18 features. Ten-fold cross-validation was explored to find their best configuration parameters. These methods are DTs, LR, and SVMs with RBF kernels or linear kernels, Gaussian-naive Bayes, and Bernoulli-naive Bayes. The well-trained models were then used to predict phage-host interactions, and the phage-host pairs discriminated by at least four models with probability greater than 0.8 were returned.

## Results

### Phage-host interaction datasets

In total, 1,751 phages and their 193 bacterial host species collected from two previous studies were split into two mutually exclusive sets for model training and external validation. A total of 817 phages and 143 annotated bacterial hosts at the species level (including 5,482 bacterial genome sequences) were used as the positive training set. The remaining 936 phages and their 110 host species (including 4,666 bacterial genomes) never used during the modeling phases were used as the positive external validation dataset. The negative training and validation datasets were built artificially by matching phages with bacteria from a different species other than their known host in a degree-preserving manner (using edge swaps but only for uniquely connecting pairs).

### Creation of custom databases

#### <u>P</u>hage <u>g</u>enome and <u>p</u>rotein <u>d</u>atabase (PGPD)

The phage genome database contained 10,463 complete phage genome sequences collected from Millardlab (http://millardlab.org/bioinformatics/bacteriophage-genomes/), which were extracted from the GenBank database on May 2019. ORFs were annotated on these phage genomes using FragGeneScan ^25^. Protein sequences shorter than 1,000 nucleotides were filtered out. Finally, 684,292 nonredundant phage protein sequences clustered using CD-HIT ^26^ were used to build the phage protein database.

#### <u>B</u>acterial <u>g</u>enome and <u>p</u>rotein <u>d</u>atabase (BGPD)

The BGPD contained 13,055 completely assembled bacterial genomes and 1,107,2607 nonredundant bacterial protein sequences obtained according to the same processing steps as those used for the *PGPD*.

#### <u>S</u>equence <u>c</u>omposition <u>d</u>atabase (SCD)

The *SCD* contained *k*-mer (*k*=6) frequency and codon usage calculated for each of the 13,055 bacterial genome sequences and 10,463 phage genome sequences and homogeneous Markov models trained for each of the 13,055 bacterial genomes using the WIsH method.

#### <u>P</u>rophage <u>D</u>NA and <u>p</u>rotein <u>d</u>atabase (PDPD)

The prophage DNA database contained DNA sequences of 63,352 prophage regions identified in 9,646 bacterial genomes using Phage_Finder or DBSCAN-SWA (our in-house developed prophage detection tool). The prophage protein database contained 345,086 protein sequences predicted using FragGeneScan in these prophage regions.

#### <u>C</u>RISPR spacer <u>d</u>atabase (CSD)

A total of 65,170 CRISPR arrays were identified from 11,162 bacterial genome sequences in the *PGPD* using CRT, CRISPRFinder or PILER-CR. Additionally, 91,685 CRISPR arrays detected from 62,176 bacterial and 167 archaeal organisms were collected from the CRISPRminer database ^27^. By merging the above CRISPR arrays, 418,766 spacer sequences together with their bacterial host information were extracted to build the *CSD*.

#### <u>P</u>rotein-<u>p</u>rotein interaction (PPI) <u>d</u>atabase (PPID)

To extract the PPIs that constitute the base for phage-host prediction, i) PPIs between bacterial and phage proteins were inferred through checking the PPIs of their homologous proteins in the IntAct Molecular Interaction Database (https://www.ebi.ac.uk/intact/) and ii) comparing the frequencies at which these PPIs occur in the positive and negative training sets (occur more than twice in the positive compared with negative training set). Finally, 912 nonredundant PPIs remained and were considered to be correlated with phage-host interactions. In the same way, 318 nonredundant Pfam domain-domain interactions (DDIs) were selected and used for further evaluation of phage-host interactions.

### Evaluation of the predictive power of diverse *in silico* signals

We designed five categories of features to represent diverse *in silico* PHISs that contribute to the prediction of phage-host interactions. First, since temperate phages are ubiquitous in nature, with nearly half of sequenced bacteria bearing lysogens ^19^, we could link phages with their bacterial hosts through identifying the integrated prophages and comparing them with the phage genomes. Thus, we introduced the prophage-related features (PROP_num_, PROP_idn_, and PROP_cov_) defined to evaluate the similarity between integrated prophage(s) and phage genomes based on homologous protein alignment by DIAMOND BLASTP and nucleotide alignment by BLASTN. Second, given the phenomenon that the phages ameliorate their nucleotide composition toward that of their bacterial hosts, we next introduced sequence composition features (S_2_* similarity, WIsH score, and codon usage score) to reflect highly similar patterns in codon usage or short nucleotide words (*k*-mers) shared between some phages and their hosts. Third, as CRISPR-Cas systems have been found in ∼45% of bacterial genomes ^20^, and approximately ∼7% of all detectable spacers could match known sequences, the majority of which originate from phage genomes ^21^, we introduced CRISPR features (CRISPR_num_, CRISPR_idn_, and CRISPR_cov_) to identify past infections between a phage and its hosts. Fourth, we incorporated genetic homology features (ALN_hpc_, ALN_idn_, and ALN_cov_) to represent genetic homolog sequences that were acquired during a past infection event. Finally, domain-domain interaction (DDI) scores (DDI_num_, DDI_bap_, and DDI_php_) and PPI scores (PPI_num_, PPI_bap_, PPI_php_) were combined to evaluate interactions between the proteins of phage and their bacterial hosts.

To assess the discriminatory power of each of the eighteen PHIS features, a one-sided t-test was used to determine the difference between the mean score of each PHIS feature in the positive and negative phage-host pairs from a training set containing 817 phages and 143 host bacterial species. Our analysis revealed that all features from four of the five categories showed extraordinary discriminating abilities, except for PPI-related features, which have acceptable ability (**Fig. 2)**. First, for sequence composition analysis, the S_2_* similarity score is used to evaluate the similarity of the oligonucleotide frequency pattern of a pair of phage-host genome sequences. Positive phage-host pairs have significantly higher (*p*-value=3.06e-125, one-sided t-test) S_2_* similarity scores than negative pairs. Second, the WIsH score computes the log-likelihood of a phage genome coming from a bacterial host genome based on the Markov chain model and has significantly different medians (p-value=3.76e-91, one-sided t-test) between the positive and the negative pairs, with the positive pairs showing more codon usage similarity than the negative pairs (p-value= 2.91e-103, one-sided t-test) (**Fig. 2a**). For each of the tested phages in the positive training set or validation set, we predicted the microbe with the most similar *6*-mer, WIsH score or codon usage profile as its correct bacterial host species. We could predict correct hosts for 41.25%, 38.31% or 16.4% of phages in the training set and 29.49%, 31.41% or 17.84% for the validation set, respectively. In terms of the three prophage-related features, they showed significant discriminant power between the positive and negative pairs (*p*-value=4.45e-58, 1.854e-112 and 1.8e-56, one-sided t-test) (**Fig. 2b**). Microbes with the integrated prophages identified by BLASTN (identity⩾0.7, accumulated prophage coverage⩾0.1) or BLASTP (homology protein percentage of prophage ⩾10%) search for the query phage were the correct host species for 35.5% or 38.19% of the 817 phage, respectively, in the training set and 15.17% or 19.66% in the validation set. Third, all CRISPR scores were significantly higher for the positive phage-host pairs than the negative ones (*p*-value=1.778e-30, 1.766e-52 and 7.889e-53, one-sided t-test) (**Fig. 2c**). Microbes with strict CRISPR spacer hits (mismatch≤2, coverage≥95%) were the correct host for 34.27% of the 817 phages in the training set and 15.81% of the 936 phages in the validation set. Fourth, for genetic homology feature, positive and negative pairs were significantly different based on homologous comparison between phage and bacterial nucleotide and protein sequences (*p*-value=4.141e-62, 4.560e-82. 2.239e-67, one-sided t-test) (**Fig. 2d**). Significant hits of each genetic homology feature were the correct host species for 37.21% and 40.02% of the phages with BLASTN (identity≥0.7, accumulated phage coverage≥0.1) and BLASTP (homology protein percentage of phage genome ≥10%), respectively, in the training set and 14.21% and 20.41% of the 936 phages in the validation set. In contrast, the PPI-related features could not provide good discriminative ability (**Fig. 2e, 2f**). The discriminating ability of these features was also validated with AUC values. Likewise, except for DDI-related features that suggested weak discrimination, the other features could achieve excellent discriminating ability with AUC values (∼0.792) (see **Supplementary Table S1** for details of the statistics).

**Fig. 2.**
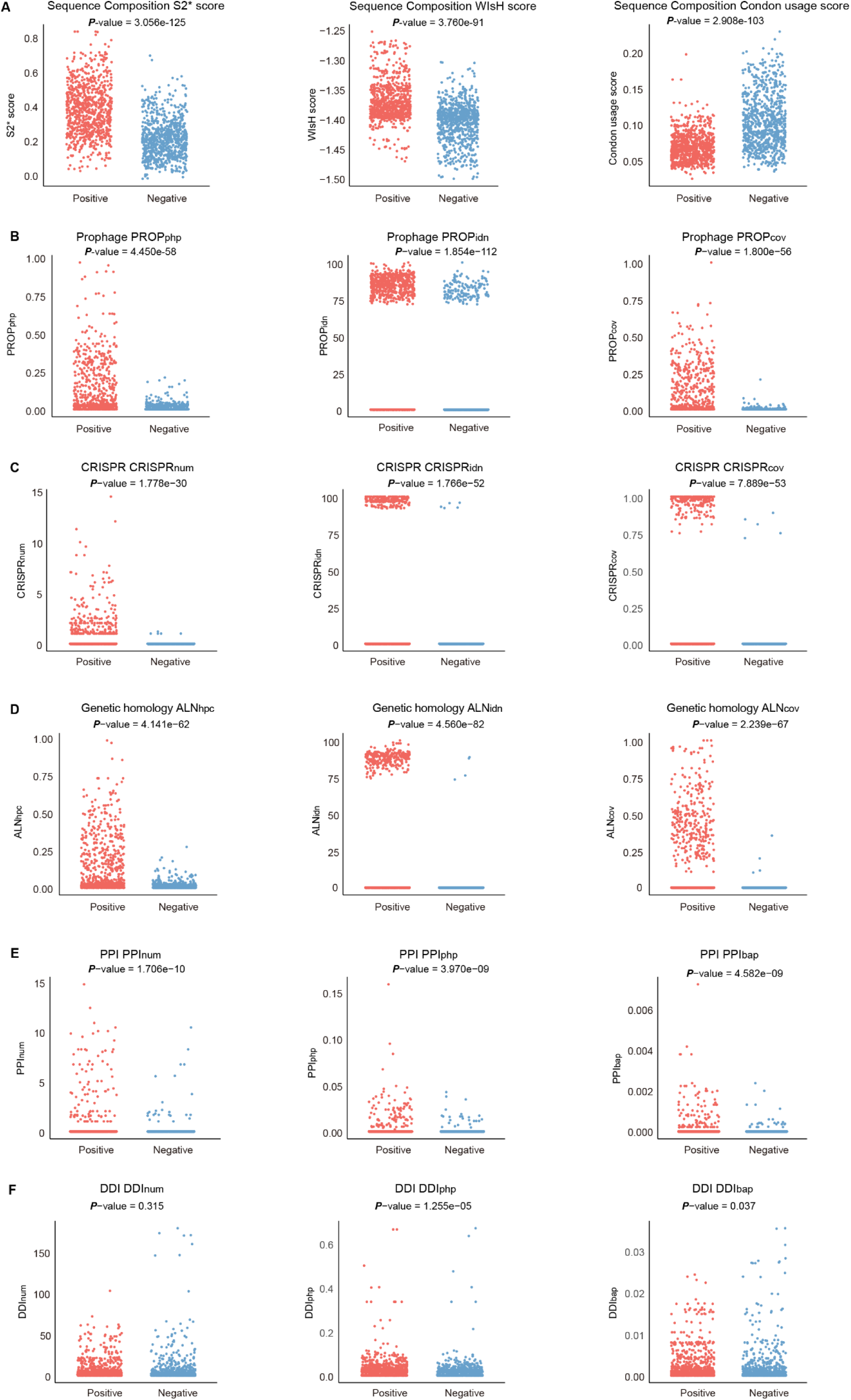
Distributions of 18 PHIS feature values in 817 interacting phage-host pairs and noninteracting phage-host pairs. (a) Jitter scatter plots of sequence composition feature values, including S2* score, WIsH score and codon usage score. (b) Jitter scatter plots of phage-related feature values, including PROP_php_, PROP_idn_ and PROP_cov_. (c) Jitter scatter plots of CRISPR-related feature values, including CRISPR_num_, CRISPR_idn_ and CRISPR_cov_. (d) Jitter scatter plots of genetic homology feature values, including ALN_hpc_, ALN_idn_, and ALN_cov_. (e-f) Jitter scatter plots of protein-protein interaction (PPI)-based feature values. (e) PPI (homology search)-based features, including PPI_num_, PPI_php_, PPI_bap_, and (f) DDI-based features, including DDI_num_, DDI_php_, DDI_bap._

### Machine learning models for phage-host interactions prediction

We observed that a single PHIS category could identify only a limited number of positive interactions, 16.4%∼41.25% for the training set **(Fig. 3a)**, while combining multiple categories of PHIS features could cover many more known interactions at the species level, approximately 70.13% of the training set. Overall, all five categories of PHIS features could together increase the possibility of capturing many additional interacting signals derived from different known mechanisms (see Materials and Methods for the details of the definitions). We attribute this phenomenon to the possibility that different types of PHISs may reflect distinct interacting mechanisms that are requisitioned by different phage-host sub-populations.

**Fig. 3.**
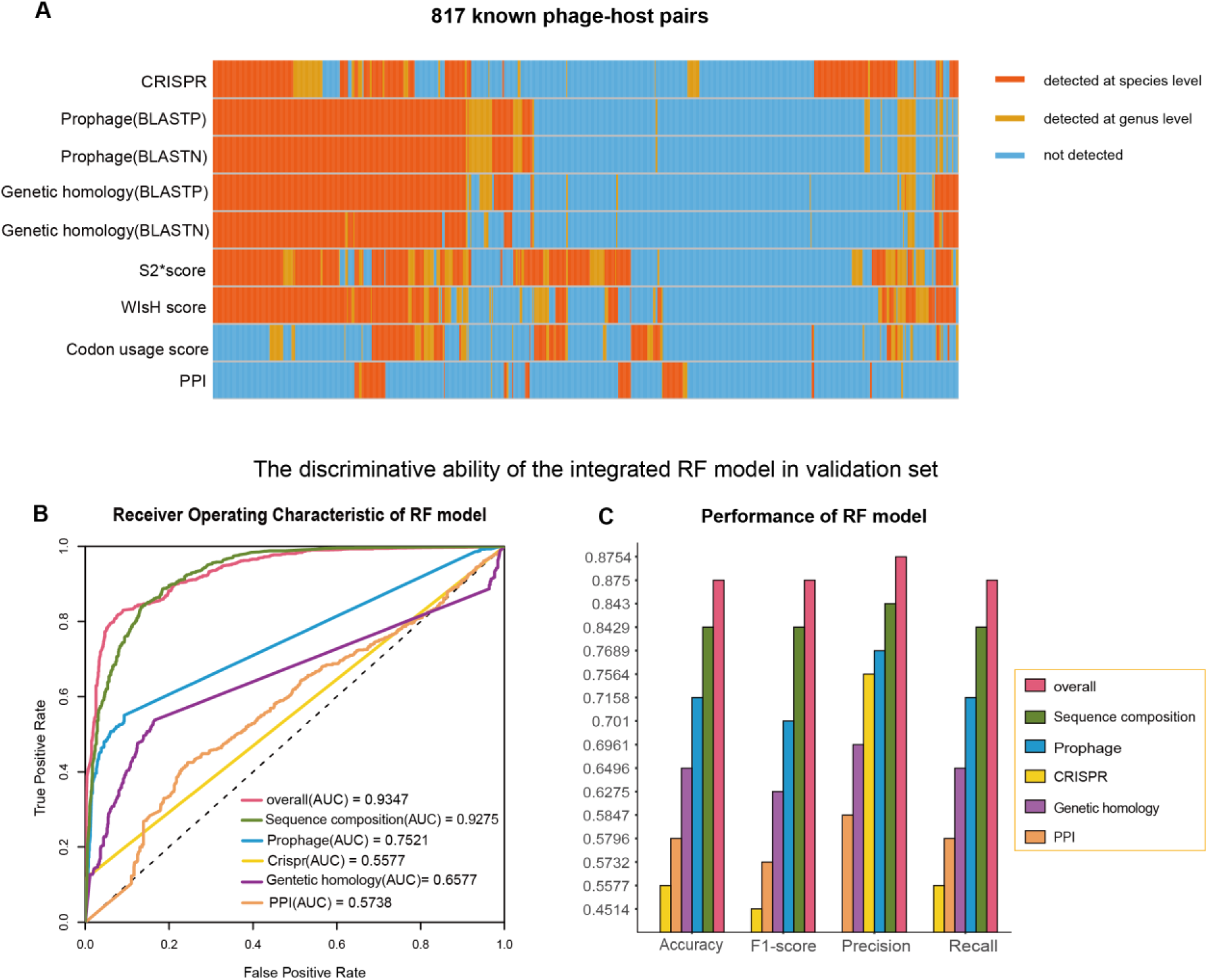
The predictive power of single PHISs and our integrated RF model. (a) Heatmap showing whether a known phage-host pair is validated by each type of *in silico* PHIS detected using different tools. Orange or yellow denotes that a phage-host pair is validated by the approach at the species level or at the genus level, respectively. Blue denotes not validated by an approach. (b) ROC curve indicating that the integrated model combining all PHISs performed better than a single model trained for a single PHIS, with the best AUC value of 0.9347. (c) Histograms displaying the performance of the integrated RF model in the validation set compared with that of other single models by four evaluation indexes, namely, accuracy, F1-score, precision and recall. The bar chart indicates that the integrated model performs better than each PHIS category, with the best accuracy of 0.875, best F1-score of 0.87496, best precision of 0.8754 and best recall of 0.875.

Therefore, based on the calculation of the above 18 *in silico* PHIS features, we then carried out machine-learning modeling to systematically integrate various categories of PHISs to predict and evaluate phage-host interactions. Seven machine-learning models, namely, random forest (RF), decision trees (DTs), logistic regression (LR), and support vector machines (SVMs) with radial basis function (RBF) kernel or linear kernel, Gaussian-naive Bayes, and Bernoulli-naive Bayes, were applied on the training dataset containing 817 phages and 143 host bacterial species with the above eighteen features. Ten-fold cross-validation on the training set was performed to find their best configuration parameters. We then applied these trained models to the positive and negative validation sets containing 936 phages infecting 110 host species independent from the training set. The overall prediction framework is demonstrated in **Fig. 1**.

To further prove that the machine learning model integrating all PHIS categories perform better than nonintegrated models, we also trained seven machine-learning models using each individual PHIS category, and the corresponding receiver operating characteristics (ROCs) using RF models were plotted. The area under the ROC curve (AUC), which measures the discriminative ability between positive and negative pairs in the validation set, was 0.574∼0.928 for each PHIS category (sequence composition, CRISPR, prophage, genetic homology and PPIs) and 0.935 for our integrated model (**Fig. 3b**). In addition, by calculating the four general evaluation indexes, including accuracy, F1-score, precision and recall, we showed that our integrated model performed much better than each single PHIS category, reaching 0.875 in all these indexes (**Fig. 3c**). In short, our approach which integrates five PHIS categories by machine-learning models exhibited dramatic predictive power for phage-host interactions.

#### Insights from analyses using PHISDetector

##### Case Study: Identification of hosts of viral contigs in a metagenomic study using a stand-alone version of PHISDetector

As a large number of new viral genomes or sequence fragments unveiled by viral metagenomics, predicting the microbial hosts for metagenomic phage contigs is one of the most fundamental challenges in understanding the ecological roles of phages ^22^. Here, we provided a stand-alone version of PHISDetector to expand our framework for large viral metagenomics dataset analysis. Users can submit high-throughput sequencing-derived phage sequences as the input, and the bacterial hosts of these phages will be returned.

We tested a set of 125,842 metagenomic viral contigs (mVCs) from 3,042 geographically diverse samples ^23^ and predicted their bacterial hosts using PHISDetector. First, using criterion 1 (Fig. 1a), we could predict bacterial hosts of 13,304 (10.57%), 2,221 (1.76%) and 276 (0.22%) mVCs by matching CRISPR spacers, genetic homology of bacterial genomes and microbial prophages with mVCs. Second, using criterion 2 (Fig. 1a), 111,058 (59.13%) mVCs were retained for further evaluation by the phage-host interaction evaluation (PHIE) module (Fig. 1b) (see Materials and Methods for the detailed description). Finally, 65,664 mVCs were returned with predicted hosts at the genus level, supported by at least two trained machine-learning models with probability ≥ 0.8. Compared with the original study in which only 9,607 (7.7%) of the mVCs were predicted mainly through CRISPR spacers and transfer RNA matches, PHISDetector annotated 68,634 (54.54%) of the mVCs, and the predicted hosts at the genus level matched the previous annotation in 62.34% of the cases (**Fig. 4**). In brief, it is convenient and fruitful to predict bacterial hosts for virome contigs using the stand-alone version of PHISDetector.

**Fig. 4.**
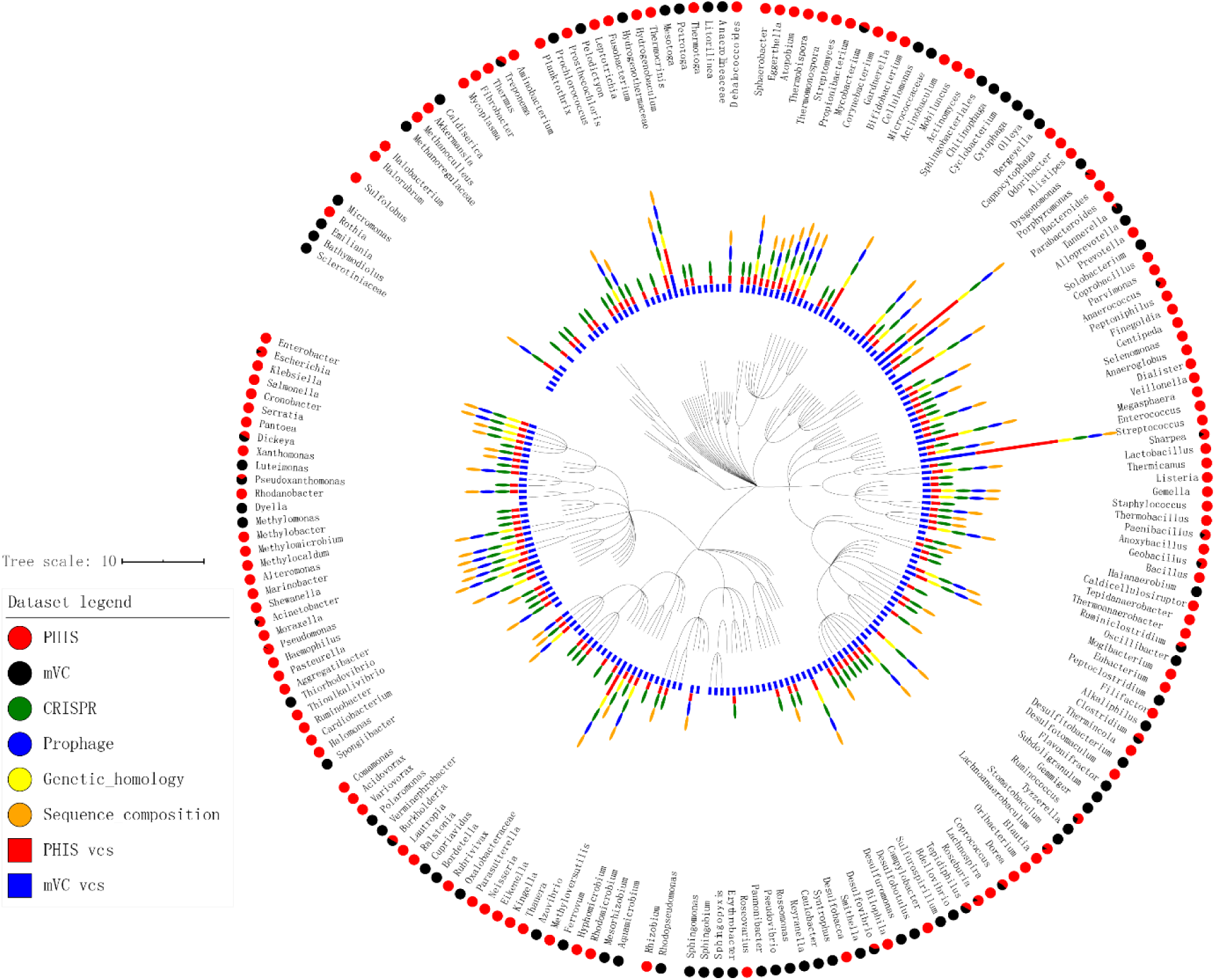
Prediction power of PHISDetector for metagenomics mVCs. Phylogenetic distribution of bacterial hosts. In total, 205 genera are shown in a phylogenetic tree. The innermost circle (blue rectangles) shows the number of mVCs assigned to this genus by Paez-Espino *et al*. The adjacent second circle (red rectangles) shows the number of mVCs assigned to this genus by PHISDetector. In the outer circles, genera are marked for detecting signals: CRISPR (green ovals), prophage (blue ovals), genetic homology (yellow ovals), and sequence composition (orange ovals). In the outermost circle, the pie charts indicate the host prediction consistency at the genus level between PHISDetector and Paez-Espino *et al*. Red indicates the fraction of mCVs assigned to this genus by Paez-Espino *et al* that is also correctly predicted by PHISDetector. Black indicates the fraction of mCVs assigned to this genus by Paez-Espino *et al* but not predicted as the same host genus by PHISDetector.

##### Case Study: Making predictions and annotations using the PHISDetector webserver

The PHISDetector webserver receives bacterial or virus genomic sequences in GenBank or FASTA format as input and provides well-designed result visualizations and data tables with details to download. Generically, for a FASTA input file, open reading frames (ORFs) will be first predicted on the input genome using FragGeneScan, while for a GenBank (GBK) file, DNA sequences and ORF amino acid sequences of the genome will be extracted directly from the input GBK file (**Supplementary Figure 1**). The PHISDetector webserver supports three types of analysis: 1) ***Evaluate interacting probability for a pair of phage and prokaryotic genomes***. If a pair of phage-microbe genome sequences has been submitted, diverse *in silico* PHISs (18 features) will be detected to characterize the interaction (**Supplementary Figure 1**, lower panel). Then, the PHIE module will be applied to indicate the possibility of the interaction. 2) ***Predict the infecting phage for a query prokaryotic genome***. If a bacterial sequence has been submitted (**Supplementary Figure 1**, upper left), the ORFs, prophage regions and CRISPR arrays will be initially detected. Then, all 18 features will be calculated between the input bacterial sequence and each of the 10,463 phages in our custom phage genome and protein database (***PGPD***) (see Materials and Methods for the detailed description). The phages passing criterion 1 will be directly returned as the infecting phages. The remaining phages passing criterion 2 will be considered candidate phages and further evaluated using the PHIE module. Next, the phages passing criterion 1 or approved by the PHIE module will be returned as the final list of infecting phages. 3) ***Predict the bacterial host for a query phage genome***. If a phage sequence has been submitted (Supplementary Figure 1, upper right), all 18 features will be calculated between the input phage sequence and each of the 13,055 bacterial genomes in our custom bacterial genome and protein database (***BGPD***) (see Materials and Methods for the detailed description). As above, the bacteria passing criterion 1 or approved by the PHIE module will be returned as the final list of bacterial hosts.

We illustrated the output results using the prediction for infecting phages of *Staphylococcus aureus* subsp. *aureus* JH1 (NC_009632), the bacterial hosts of *Staphylococcus* phage 47 (NC_007054), and the characterization of this phage-host interaction. The PHISDetector webserver first displays the predicted bacterial hosts and diverse *in silico* phage-host interaction signals in the form of interacting tables, which are easy to query, sort and download. For each phage-host pair in the consensus table, powerful, exquisite and beautiful interactive visualization graphics were provided for every *in silico* PHIS. The interactive DataTables are used to display the prediction results with details for CRISPR, prophage, genetic homology and PPI. Furthermore, several kinds of interactive graphics are provided for users to better understand the prediction results. For CRISPR, the table shows the predicted hosts, and the detailed spacer targets yield the host *<u>Staphylococcus argenteus</u>*<u>strain 3688STDY6125086</u> (<u>NZ_FQRJ01000002</u>) in Fig. 5 according to the spacer that matched the query phage with the identity of 0.94 and coverage of 0.972 by performing a BLASTN search against our underlying ***CRISPR spacer database (CSD)***. For prophage, an interactive circular genome viewer is used to illustrate the prophage regions of the bacterial genome with the hit prophages highlighted in red and equipped with captions next to it, and DataTables to display the detailed hits between each prophage region and *Staphylococcus* phage 47 in Fig. 5 by performing a BLASTP and BLASTN search against the phage, yielding three hit records with the best prophage homology percentage, identity and coverage of the prophage greater than 0.7, 89% and 0.84. For Genetic_homology, circular genome viewers are used to display the homologous proteins and matched regions of *Staphylococcus* phage 47 in Fig. 5 by performing a BLASTP and BLASTN search against *Staphylococcus aureus* subsp. *aureus* JH1, respectively, with the color shade representing the degree of matching as well as tables to show the detailed matches. For sequence composition, density curves are plotted to comparatively show the sequence composition similarity between the phage-host pair based on the background density distribution of reference datasets (817 known phage-host pairs) and corresponding blue reference lines, indicating the high similarity of sequence composition between the phage-host pair in Fig. 5. For PPIs, interactive bipartite networks present the PPIs between the phage and bacterial proteins. Concerning prediction for infecting phages of *Staphylococcus aureus* subsp. *aureus* JH1 (NC_009632) and the characterization of this phage-host interaction, the visualization graphics are similar to those above (**Fig. 5** and **Supplementary Figure 1)**. In addition, PHISDetector also provides seven independent analysis modules, namely, oligonucleotide profile analysis, CRISPR analysis, phage analysis, similarity analysis, cooccurrence analysis, specialty gene check, and PPI analysis, to provide a flexible and convenient one-stop web service of phage-host interaction-related analysis (**Supplementary Figure 2)**.

**Fig. 5.**
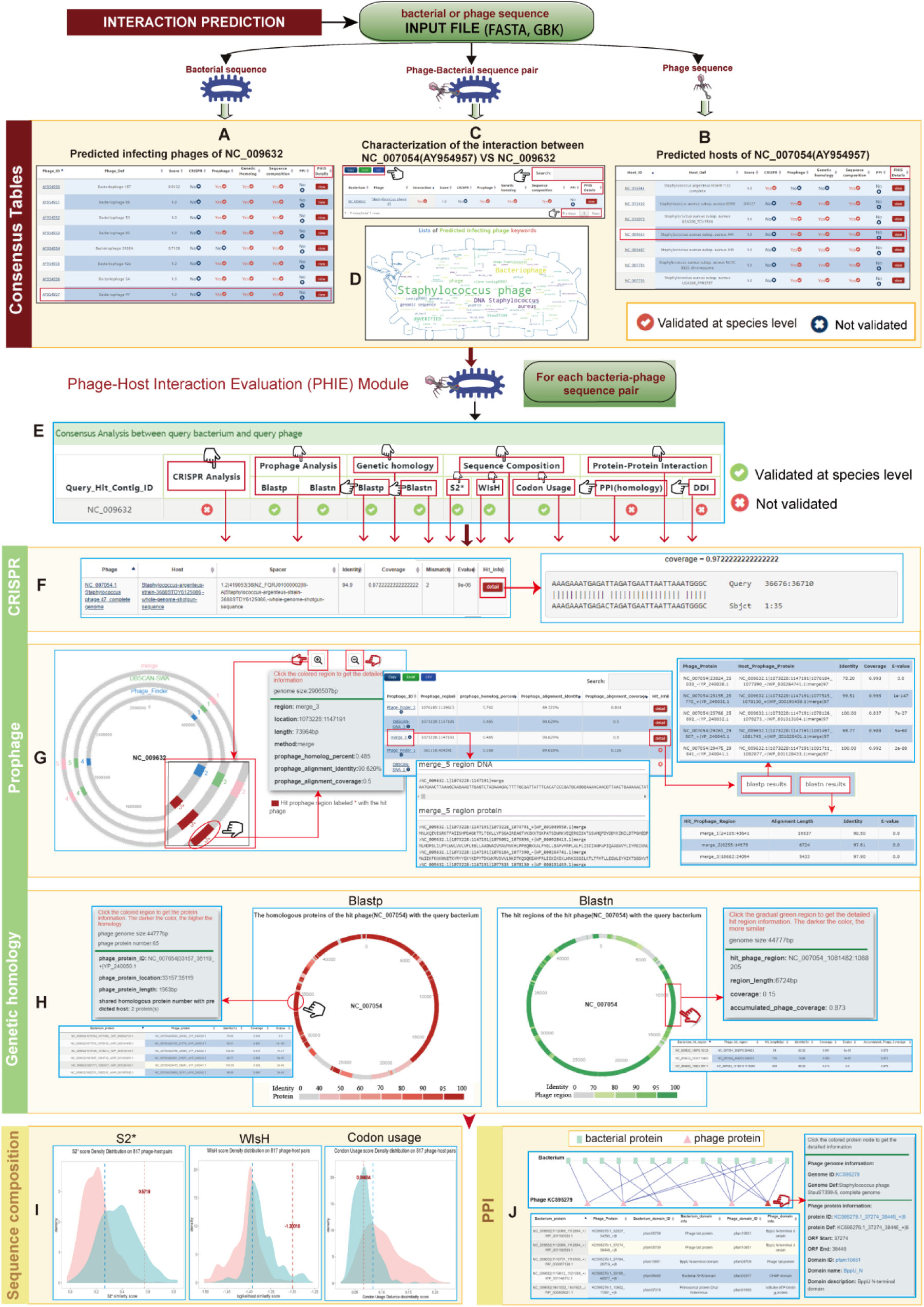
Illustrations for making predictions and annotations using the PHISDetector webserver. (a-c) Predicted infecting phages of *Staphylococcus aureus* subsp. *aureus* JH1 (NC_009632), predicted hosts of *Staphylococcus* phage 47 (NC_007054 or AY954957) and the characterization of this phage-host interaction with detected PHIS signals in each bacterium-phage pair. (d) Predicted phage keywords. (e) Consensus *in silico* detection of each PHIS in this interaction (NC_009632 vs NC_007054). (f) An example of CRISPR visualizations of predicting the hosts of *Staphylococcus* phage 47, including the predicted host table and detailed spacer targets, yields the host *<u>Staphylococcus argenteus</u>*<u>strain 3688STDY6125086</u> (<u>NZ_FQRJ01000002</u>). (g) An example of phage visualizations including a circular genome viewer to illustrate the prophage regions of *Staphylococcus aureus* subsp. *aureus* JH1 with the hit prophages highlighted in red and equipped with captions next to it and DataTables to display the detailed hits between each prophage region and *Staphylococcus* phage 47, showing the hit proteins and regions in tables by clicking the detail button. (h) An example of genetic homology visualizations consisting of circular genome viewers to display the homologous proteins and matched regions of *Staphylococcus* phage 47, with the color shade representing the degree of matching and captions attached to the gray frame by clicking the region, as well as tables to show the detailed matches. (i) Density curves are plotted for comparative evaluations of the sequence composition similarity between the phage-host pair (NC_009632 vs NC_007054) in terms of S2*, WIsH and codon usage scores (red line) based on the density distribution of reference datasets (817 known phage-host pairs) and corresponding blue reference lines. (j) Interactive bipartite network and tables to present the PPIs between *Staphylococcus aureus* subsp. *aureus* JH1 and the predicted infecting phage *Staphylococcus* phage StauST398-5 (<u>KC595279</u>).

## Conclusions and discussion

Phage-host interaction is one of the essential questions in microbial environments. Despite rapidly expanding knowledge of the virosphere from culture-independent metagenomic sequencing surveys, only a fraction of its diversity has been described using classic cultivation techniques. Sequence-based characterization of uncultured viral diversity requires major overhauls in the current viral classification schemes and has greatly improved methods to link uncultured viruses to their hosts, which are almost entirely unknown currently ^24^. Previous studies have indicated that molecular and ecological coevolutionary processes of phages and bacteria leave signals in their genomic sequences for tracing their interactions. Several computational tools have been developed to detect various signals, such as integrated prophages, oligonucleotide frequency patterns, and CRISPR spacer sequences. In this paper, we applied an integrated approach to develop PHISDetector for predicting phage-host interactions. Compared with prior tools, the PHISDetector pipeline described here is uniquely comprehensive because it integrates various types of PHISs reflecting possible phage-microbe interacting mechanisms into one tool and adds valuable novel functionalities. Consequently, PHISDetector can predict additional interactions that cannot be detected if using only one single category of signals and calculate the possibility of a novel phage and microbe pair with extreme precision. Users can choose the web server or stand-alone version flexibly according to their research and resources. Both provide well-designed, interactive visualization outputs for improved result interpretation and illustration for further analysis.

Our prophage analysis module combined two popular programs, Phage_Finder and VirSorter, and our in-house developed tool, DBSCAN algorithm [31] combined with sliding window algorithm (SWA) (DBSCAN-SWA). DBSCAN-SWA presents the best detection power based on the analysis using a controlled dataset including 184 manually annotated prophages, with a detection rate of 85%, which is greater than that of Phage_Finder (63%) or VirSorter (74%). If combining all three methods (provided as a “merge” function in the prophage analysis module), 92% of the reference prophages could be detected. We also added a prophage annotation step to indicate the possible integrated phages of the predicted regions. Our CRISPR analysis module facilitates two-way analysis. If a phage genome is submitted, it will be compared with our in-house collected spacer database (***CSD***, including 418,766 spacer sequences from 63,182 bacterial and archaeal genomes) to quickly detect CRISPR-targeting associations between the input phage sequence and microbial genomes in NCBI. If a bacterial genome is received, PHISDetector will detect the CRISPR spacer automatically and compare against an in-house collected phage genome database (*PGPD*) to find the target phage. If a bacterium-phage pair is received, PHISDetector will detect the CRISPR spacer automatically in the bacterial genome sequence and compare against the query phage genome to predict the CRISPR-targeting association. The sequence composition analysis module supports VirHostMatcher, WIsH, and codon usage, which are complementary to each other because VirHostMatcher may be more suitable for complete genomes, while WIsH (for virus contigs shorter than 10 kb) and codon usage distance can be detected in both complete and incomplete genomes. The genetic homology module detects the exact matches between any phage-host pair regions of genetic homology and provides well-designed visualizations to display the degree of matching between the phage-host pair by generally colored circular genome viewers. In the coabundance analysis module, we used the CoNet program to infer a viral and bacterial cooccurrence network. As a plug-in of Cytoscape, we adapted CoNet to a web version to better aid biologists without computational background to use and adjust parameters. We also provided a functional module for the detection of PPIs and DDIs between a pair of phage-host genomes to better understand their interplay at the protein level. In addition, to assist the characterization of phage genomes for therapeutic applications, we introduced a specialty gene check module to detect virulence factors (VFs) and antibiotic resistance genes (ARGs). Finally, a consensus analysis using machine-learning models is performed to indicate the possible integrity of the predicted interactions and interplay among different PHISs. Based on PHIS detections for the training set consisting of 817 known phage-host interactions, more than 85% of the phage hosts were correctly identified at the species level by combining various approaches. Furthermore, the integrated RF model trained based on the training set attained the best performance, with an <u>AUC value of 0.935</u> and accuracy of 0.875 for the validation set (936 known phage-host pairs). Therefore, PHISDetector can predict interactions that could not have been detected if using merely a single category of signals and calculate the possibility of novel phage-microbe pairs with extreme precision.

In summary, PHISDetector is a tool that can detect diverse PHISs and reflect various interaction patterns or mechanisms so that users can easily compare the results from different methods and better understand phage-bacteria coevolution. PHISDetector offers well-designed and interactive visualization outputs for improved result interpretation. Users can choose the web server or stand-alone version to fit their research needs. PHISDetector will continue to develop to incorporate additional *in silico* phage-host signals and evaluate the consistency or association of various signals upon extensive analysis of large data sets. We hope that PHISDetector can promote research on the roles of phage-host interactions from ecological and evolutionary perspectives, facilitate our understanding of their roles in human health and disease, and accelerate the development of novel therapeutic strategies, such as modulating specific microbes in a microbial community and treating multidrug-resistant infections.

## Methods

### Calculation of PHIS features

We considered that diverse *PHIS* features in bacterial and phage genomes potentially impinge on host range determination, including sequence composition similarity, CRISPR targeting, prophages, regions of genetic homology, and PPIs or DDIs. We constructed eighteen features belonging to five categories in our framework. A detailed definition of individual features is provided in **Supplementary Table S2**. (i) ***Sequence composition-related features*** (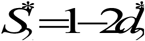, WIsH score and codon usage score); 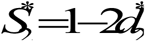 and WIsH score are calculated based on estimated *k*-mer frequencies ^10 11^. The codon usage score is evaluated as the dissimilarity in codon usage profiles of phage and bacterial coding regions. (ii) ***CRISPR-related features*** (CRISPR_num_, CRISPR_idn_, and CRISPR_cov_) represent the number of shared CRISPR spacers, the best identity and coverage over all the hits between the host spacers and the phage genome, respectively. (iii) ***Prophage-related features*** (PROP_php_, PROP_idn_, and PROP_cov_) are defined to evaluate the similarity between the prophage region(s) on bacterial and phage genomes based on homologous protein comparison and nucleotide similarity. (iv) ***Genetic homology features*** (ALN_hpc_, ALN_idn_, and ALN_cov_) represent the exact matches between phage and bacterial genome regions of genetic homology. (v) ***PPI- or DDI-based features*** (PPI_num_, PPI_bap_, PPI_php_, DDI_num_, DDI_bap_, and DDI_php_); six features were calculated to evaluate the interacting potential for every phage-host pair based on the interactions between their encoded proteins: (i) the number of PPIs or DDIs between the bacterial proteins and phage proteins (PPI_num_ and DDI_num_); (ii) the proportion of bacterial proteins involved in PPIs or DDIs (PPI_bap_ and DDI_bap_); and (iii) the proportion of phage proteins involved in PPIs or DDIs (PPI_php_ or DDI_php_).

### General workflow

PHISDetector combines two stages. In stage I, phage-host pairs with high reliability were detected using criterion 1 (**Supplementary Table S3**) and returned directly as final predicted results. In stage II, phage-host pairs with potential PHISs based on criterion 2 (**Supplementary Table S3**) were retained and further evaluated using trained machine-learning models implemented in the PHIE module (**Fig**. 1a).

#### Phage-host interaction evaluation (PHIE) module

Seven machine-learning models were trained on the training dataset with the above 18 features. Ten-fold cross-validation was explored to find their best configuration parameters. These methods are DTs, LR, SVMs with RBF kernels or linear kernels, Gaussian-naive Bayes, and Bernoulli-naive Bayes. The well-trained models were then used to predict phage-host interactions based on 18 PHIS features (**Supplementary Table S2**), and the phage-host pairs discriminated by at least four models with probability at least 0.8 were returned (**Fig. 1b**).

#### Evaluation methods

One-sided t-tests were used to examine whether the signal scores were significantly different between positive and negative phage-host pairs. ROC curves were used to assess the power of predictive signals by plotting the false positive rate (100-specificity) versus the true positive rate (sensitivity) according to the change in threshold for each signal feature. The AUC is a measure of the ability of the model to rank true interactions higher than noninteractions independent of the prediction score threshold. The values of sensitivity (true positive rate) and specificity (true negative rate) were used as accuracy metrics for users to better assess the prediction results. All analyses were carried out using the Python package ‘scikit-learn’. Feature contributions to the models were output as feature importance.

### Integrated analysis tools

PHISDetector is composed of three types of analysis modules that allow (i) identifying diverse *in silico* PHISs, including oligonucleotide profile analysis, CRISPR analysis, prophage analysis, similarity analysis and cooccurrence/coabundance analysis; (ii) checking specialty genes, including VFs and ARGs; and (iii) detecting PPIs between a pair of phage and bacterial genomes. These integrated tools can be accessed via http://www.microbiome-bigdata.com/PHISDetector/index/tools/.

#### CRISPR analysis

The CRISPR spacer sequences are computationally identifiable sequence signatures of previous phage-host infections. In this module, three scenarios of analysis are supported. (i) Users can provide their input either as spacer sequences in (multi-)FASTA format; as CRISPRFinder [20], PILER-CR [29] or Seq2CRISPR [30] output files; or as a bacterial genome sequence for which the CRISPR spacers will be automatically identified using PILER-CR. Next, putative protospacer targets will be identified by a BLASTN search of the spacer input against the viral reference database. (ii) Users can upload viral sequences that will undergo BLASTN search against the spacer reference database. Two spacer reference databases have been built in our pipeline, including spacers predicted from complete and/or draft bacterial genomes in NCBI. The bacterial sources of the identified spacers are predicted as the potential hosts of the viral sequences. (iii) Users can check the phage-host links by CRISPR spacer-protospacer matching between the uploaded bacterial and phage sequences in (multi-)FASTA format. The spacer sequences will be predicted on the bacterial sequence using PILER-CR first and will be aligned to the phage sequences by BLASTN.

#### Prophage analysis

The prophage analysis module accepts both raw DNA sequences in FASTA format and annotated genomes in GenBank format and provides three prophage detection programs, including Phage_Finder, VirSorter, and DBSCAN-SWA (unpublished). DBSCAN-SWA implements an algorithm combining the DBSCAN algorithm [31] and SWA, referring to the theory of PHASTER, a widely used web tool for prophage prediction with no stand-alone version or source code available [4]. In addition, tRNA sites are annotated using ARAGORN [34] for raw DNA sequences and extracted for annotated sequences. Sequences of 10 upstream and downstream proteins of each cluster using integrase as the anchor were extracted to examine putative att sites using BLASTN with parameters ‘-task blastn-short –evalue 1000’. Finally, the characterization of the predictive prophage region is performed using BLASTN against the UniProt viral genome DNA sequences, and the best hitting phage organism is returned. We also used the viral UniProt TrEML reference database to annotate the predicted ORFs in the prophage region. Annotated ORFs with taxonomy information were then subjected to a voting system, and the prophage region was assigned a taxonomy based on the most abundant ORF taxonomy annotated within the prophage. The distribution of prophage-like elements detected by different methods and their size relative to the genome of their host are shown on an interactive circular genome viewer encoded using AngularPlasmid (http://angularplasmid.vixis.com). The corresponding prophage annotation is shown in the right panel when clicking on the regions.

#### Oligonucleotide profile analysis

This module predicts the bacterial host of phages by examining various oligonucleotide frequency (ONF)-based distances/dissimilarities using VirHostMatcher. For the prediction of the prokaryotic host of short viral contigs, an extra WIsH approach is provided. Note that an extra taxonomy file is required when using the VirHostMatcher approach, so we provide a tool to generate the taxonomy file for the input bacterial genomes using NCBI accession IDs.

#### Specialty gene check

As accessory genetic elements, bacteriophages play a crucial role in disseminating genes and promoting genetic diversity within bacterial populations. They can transfer genes encoding VFs such as toxins, adhesins and aggressins to promote the virulence of the host bacteria. Additionally, ARGs in bacterial chromosomes or plasmids can be mobilized by phages during the infection cycle to increase antibiotic resistance. To identify specialty genes for a pair of bacteria-phage genomes, ORFs were first predicted using FragGeneScan, and then ShortBRED [35] and Resistance Gene Identifier (RGI) v3.1.1 (https://github.com/arpcard/rgi) were used to search predicted ORFs against the Virulence Factors of Pathogenic Bacteria (VFDB) database [36] and the Comprehensive Antibiotic Resistance Database (CARD) [37], respectively. This analysis facilitates our understanding of how specialty genes are transferred between bacteria and phages.

#### Protein-protein interaction analysis

Interactions between bacteriophage proteins and bacterial proteins are important for efficient infection of the host cell. We assigned bacterial and phage genes to homologs in the UniProtKB protein database based on amino acid sequence homology via DIAMOND searches, and then the interactions between bacteriophage and bacterial proteins were inferred through checking the PPIs of their homologs in the IntAct Molecular Interaction Database (https://www.ebi.ac.uk/intact/). The interactions between bacteriophage proteins and bacterial proteins may contribute to understanding the infectious interactions between bacteria and phages.

---

**Supplementary Figure 1.**
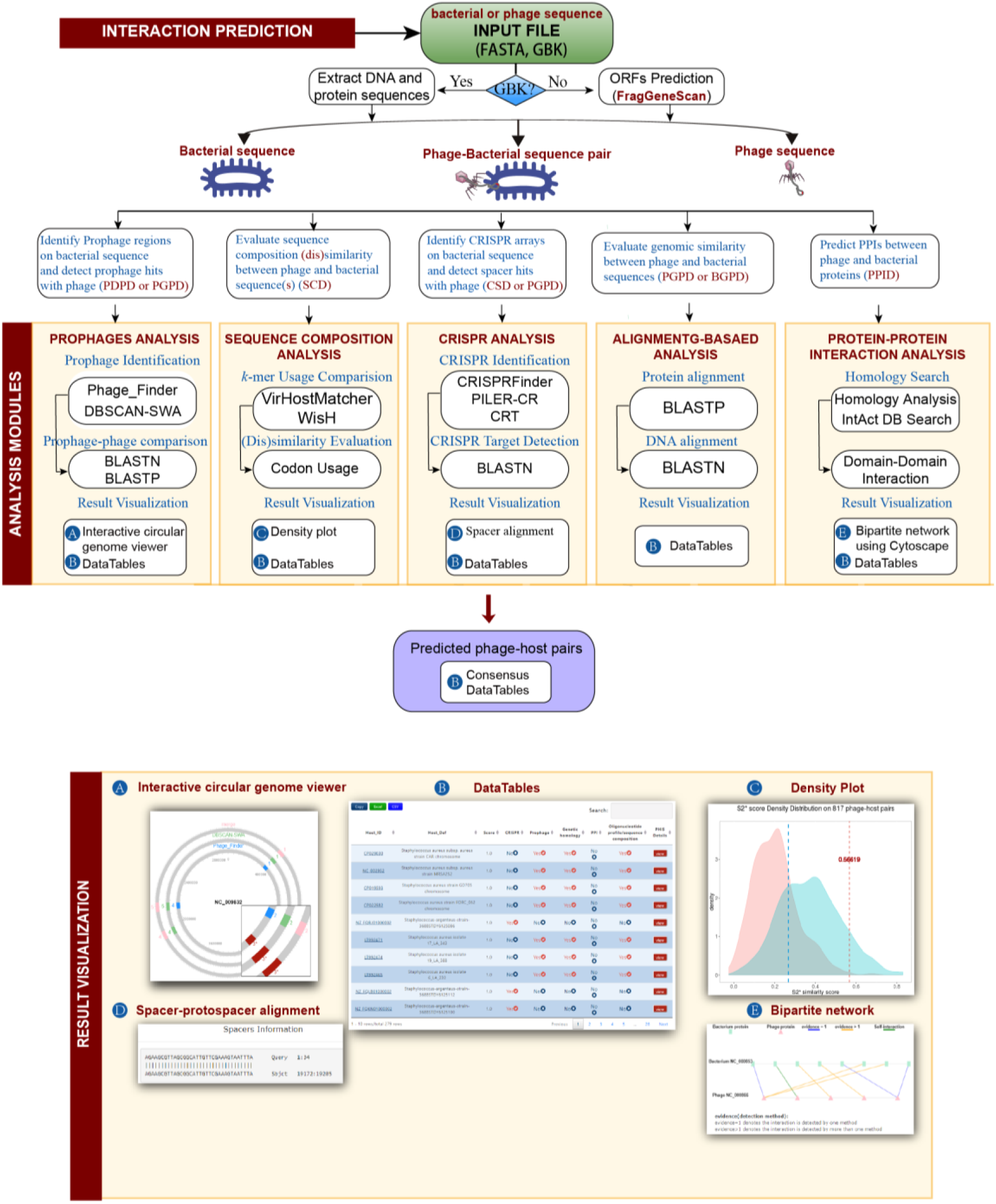
Webserver pipeline for prediction and evaluation of microbe-phage interactions by PHISDetector. The pipeline receives the FASTA sequence file and the annotation file (GenBank) of the microbe or phage genome as input for further evaluation. PHISDetector is composed of two main analysis components: (i) interaction prediction that allows predicting microbe-phage interactions using diverse methods depending on the input sequence (microbe, phage or bacterium-phage pair) (upper panel), and (ii) five underlying analysis modules (middle panel) support diverse *in silico* signal detection between any predicted bacterium-phage pair to understand their coevolution. In addition, PHISDetector uses additional modern JavaScript tools for the visualization of diverse prediction outputs.

**Supplementary Figure 2.**
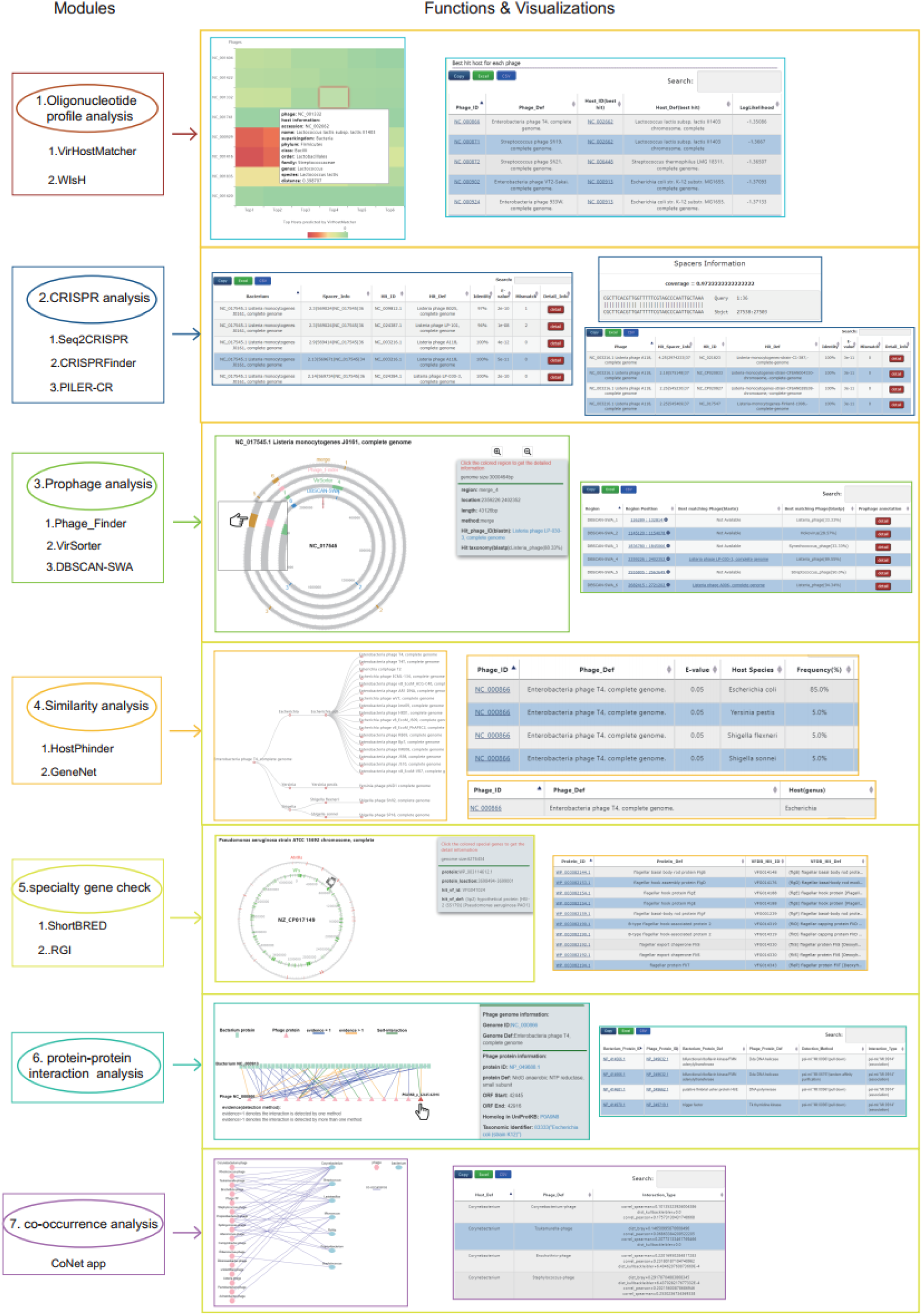

